# Comparison between conventional electrodes and ultrasound monitoring to measure TMS evoked muscle contraction

**DOI:** 10.1101/2020.02.07.938837

**Authors:** Isabella Kaczmarczyk, Vishal Rawji, John C. Rothwell, Emma Hodson-Tole, Nikhil Sharma

## Abstract

**Background:** Transcranial Magnetic stimulation (TMS) is a non-invasive cortical stimulation method that has been widely employed to explore cortical physiology in health and a range of diseases. At the core of many TMS protocols is the measurement of evoked muscle contractions using surface electromyography (sEMG). While sEMG is appropriate for many superficial muscles such as abductor pollicis brevis (ABP) and first dorsal interosseous (FDI), there are situations where the study of less accessible muscles may be of interest. Peripheral ultrasound is a non-invasive method that could provide a solution. We explore the relationship between TMS evoked sMEP and TMS evoked muscle contractions measured with muscle ultrasound. We hypothesise that in a healthy population, we expect a positive correlation between EMG and ultrasound measures.

**Methods:** In 10 participants we performed a standard TMS recruitment curve and simultaneously measured MEP and peripheral muscle ultrasound (pUS). We targeted the following muscles: biceps (BI), first dorsal interosseous (FDI), tibialis anterior (TA) and the tongue (TO).

**Results:** We report a very close relationship between the MEP and pUS contraction. Resting motor threshold (RMT) measurements and recruitment curves are consistent in sEMG and pUS. A key aspect of this work is the ability to examine clinically relevant muscles that are difficult to probe using surface EMG electrodes, such as the tongue.

**Conclusion:** We find that TMS muscle contractions can be measured with muscle ultrasound in superficial and deep muscles, enable additional, previously hard to study muscles, to be investigated. This could be valuable for allowing TMS to be used to explore a new range of muscles in disorders such as ALS. In muscles less accessible by sEMG, such as the tongue, it may be possible to use pUS as an alternative output. This may be useful in conditions such as ALS and stroke that can differentially affect the tongue.

## Introduction

Transcranial Magnetic stimulation (TMS) is a non-invasive cortical stimulation technique that has been widely employed to explore cortical physiology in health and a broad range of diseases. A TMS pulse painlessly induces descending pyramidal volleys in corticospinal tracts, the summation of which can be detected by electromyography (EMG) as motor evoked potentials (MEPs), using electrodes attached to a contralateral limb muscle. Although we presume that these fast-conducting pyramidal tracts are activated, the precise TMS-induced mechanism of action is presently undetermined (Terao and Ugawa, 2002). While there is a wide variety of TMS protocols, at the core of nearly all of them is the measurement of evoked muscle contractions using surface EMG. This defines the resting motor threshold (RMT) on which many other protocols, such as paired-pulse and repetitive pulse protocols, are based.

TMS is useful in studies involving changes in cortical excitability (Pascual-Leone et al., 1998), cerebral reorganisation, movement and neurological disorders. For example, pathologically altered corticospinal tract conduction in diseases such as MND can be attributed to a loss of corticomotorneurons, demyelination or blocked conductivity (Vucic et al., 2013; Vucic and Kiernan, 2017). TMS alone has shown a clear correlation between these conditions and increased corticospinal excitability in people living with MND, particularly in the early stages of the disease. TMS distinguishes changes in corticospinal excitability even in conditions that do not involve motor-neuronal structural damage, such as Parkinson’s disease (Leon-Sarmiento et al., 2013). This illustrates that TMS is not only sensitised to detecting the effects of a structurally impaired corticospinal tract (CST), but to the summative actions of excitatory and inhibitory corticospinal interactions, which will bring about changes in MEPs.

TMS is recorded using surface electrodes (sEMG), which have limited access to certain muscles that may be of interest in neuromuscular disorders. Primarily, surface electrodes are insufficient for monitoring MEPs in muscles of the bulbar region. For instance, monitoring the cortical physiology of the tongue may be important in MND where about 20-30% of people present primarily with bulbar involvement and many later go on to develop bulbar symptoms (Gubbay et al., 1985). Notwithstanding our particular focus on MND, there are likely many other conditions when detailed study of the tongue could be valuable. Secondly, sEMG doesn’t have the capacity to differentially register MEPs in regions where multiple muscles overlay at various depths beneath the skin. This results in an overrepresentation of signal from muscles nearer to the surface of the skin and reduced or absent signal from MEPs in deeper layers.

Muscle ultrasonography is a well-established muscle morphology visualisation tool that may overcome sEMG limitations of accessibility to certain muscles. Peripheral muscle ultrasound (pUS) can record both static and continuous muscle imagery. Firstly, it can be used to aid the diagnostic process in neuromuscular diseases by monitoring changes in muscle thickness and muscle architecture (Heckmatt et al., 1982). More importantly, pUS has the ability to register muscle movement in real-time, making it an excellent tool for detecting small intra-muscular movements, such as muscle fasciculations (Walker et al., 1990; Arts et al., 2008). Reimers et al., (1996) indicates pUS is more sensitive than sEMG examination. Recently there have been further developments in robust computational methods for detecting and analysing fasciculations using pUS (Bibbings et al., 2019). Ultrasound has become a powerful tool that can provide a broader view of typically difficult to access muscles, although it is unknown whether we can use pUS to measure TMS muscle contraction.

In this study, we aim to understand the relationship between sEMG and pUS to investigate whether we can use pUS to measure TMS muscle contraction. We expect that for difficult to access muscles, such as the tongue, pUS recordings are a more reliable measure of MEPs than traditional sEMG electrodes. We hypothesise that sEMG and pUS TMS evoked measurements will be highly correlated across a range of muscles. This will provide a foundation to use TMS-pUS to explore less accessible muscles. In a healthy population, we expect a positive correlation between EMG and pUS measures.

## Methods

### Subjects

10 right-handed subjects (5 females, mean age 23.00, SD 1.61) were recruited. The first dorsal interosseous (FDI), the biceps (BI), tibialis anterior (TA) and tongue (TO) were studied (the tongue was studied in a subset of 8 subjects). Inclusion criteria were healthy, right-handed, consenting adults of either gender. Participants were excluded from the study if they had been diagnosed with recent trauma such as fractures or structural pathologies of the right first dorsal interosseous, the right biceps, the right tibialis anterior and the tongue. Implanted metal objects or devices (cochlear implant or deep brain stimulator) in the brain or skull were prohibited. Individuals taking pro-epileptogenic medication or a history of spinal surgery were also excluded from the study. No subject had contraindications to TMS or ultrasound, which was assessed by a screening questionnaire. The study was approved by the University College London Ethics Committee.

### Experimental Setup

Standard recruitment curves were acquired in each muscle using both sEMG and pUS. Participants were subjected to 100 single pulses of TMS (10 blocks of 10 single-pulse TMS simulations at 10% intervals 10%-100% maximum stimulator output) over the hot spot corresponding to their right FDI, BI,TA and TO. Stimulation intensity was randomised and stimulation intervals included random jitter.

### TMS

TMS was carried out with a Magstim 2002 magnetic stimulator using a standard commercial figure-of-eight coil (double 70mm alpha coil). The coil was placed over the M1, tangentially to the scalp, 45° from the midsagittal line, approximately perpendicular to the central sulcus with current direction in a posterior-anterior direction. Stimulation target hotspot was determined by varying the coil direction and intensity over the motor homunculus, marking locations corresponding to maximum MEP output of the four desired muscles (BI, FDI, TA, TO).

### Electromyography (EMG)

Electromyography (EMG) was monitored by disposable electrodes (Ambu WhiteSensor 40713). Electrodes were placed bilaterally on the muscle belly of biceps (BI), first dorsal interosseous (FDI) and tibialis anterior (TA) (as seen on Fig.1). For tongue recordings, one electrode was placed on the superior body and one on the inferior body of the tongue. Electrodes were connected to an isolated amplifier system (model D360, Digitimer Ltd.), which recorded MEP data on Signal software (Cambridge Electronic Design Ltd., Version 7).

**Figure 1:**
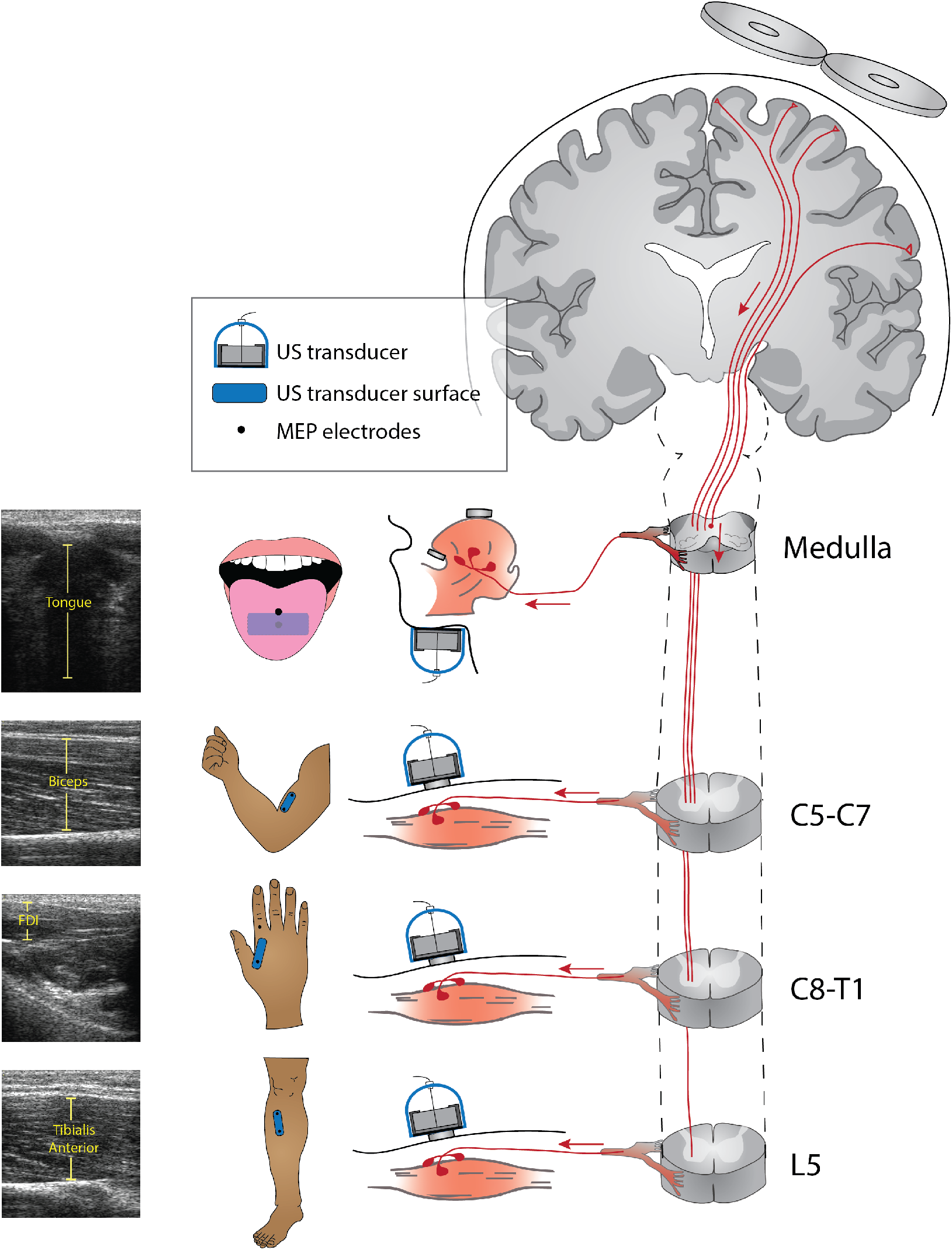
Cartoon representation of the corticomotoneuronal path from the motor cortex to the tongue, biceps, first dorsal interosseous and tibialis anterior muscles. Locations of the ultrasound probe and peripheral electrodes, which measure motor evoked potentials (MEPs), are represented. Ultrasound scans of respected muscles using the probe are shown on the left.

Resting motor threshold (RMT) on the non-dominant (right) hemisphere was established. In efforts to standardise international guidelines, an individuals’ RMT is defined as the stimulator output at which at least 5 out of 10 consecutive trials produce an MEP of at least 50μV in amplitude.

### Peripheral muscle ultrasound (pUS)

Ultrasound recordings were acquired using a Telemed LS128 ultrasound imaging system. The transducer (HL9.0/40/128Z-4) has a real-time imaging rate of~63 fps at time of recording and a frequency range set to 7MHz for skeletal muscle analysis. The system was set in B-mode with harmonics on and focal depth, focus and gain defined per muscle to optimise image quality. These settings were kept consistent between muscle types. Spectra 360 electrode gel (Parker Laboratories Inc.) was used as an acoustic medium when applying the ultrasound transducer to each of the muscles, or under the chin for tongue recordings.

### Data Processing and Analysis

#### MEP Data Processing

MEP peaks were extracted using Signal software (Cambridge Electronic Design Ltd.) and analysed offline, using R in RStudio (Version 1.1.463 RStudio, Inc.). Individual and group mean MEPs were calculated at each stimulation frequency. Pearson correlations determined the relationship between sEMG and pUS.

#### pUS Data Processing

Computational image analysis approaches were used to quantify TMS evoked twitches in recorded ultrasound image sequences. For the BI, FDI and TA a Lucas-Kanade feature tracking (Lucas and Kanade, 1981) based approach was used to capture the muscle tissue displacements evoked by TMS. The process is described in detail elsewhere (Darby et al., 2012; Harding et al., 2016). Briefly, this involved placing an evenly spaced 80 × 100 feature grid on the image. An iterative search to identify the position of each feature in the subsequent image was then completed and the total movement of all features between the two images calculated. The process was repeated for all recorded images in a sequence. The resulting total displacement values were smoothed (Lowpass butterworth filter 5 Hz cut-off) and peaks, greater than the threshold (signal mean + 0.25 × Standard deviation), identified. The time of each peak, identified from the individual frame timestamps (Miguez et al., 2017), and magnitude (pixels) was recorded for statistical analysis.

Underlying movements related to breathing and swallowing were captured in the ultrasound image sequences of the TO. These caused displacement in the feature tracking results that influenced magnitude measures of evoked muscle contractions. Therefore, for the TO a recent foreground detection based approach was used (Bibbings et al., 2019). This approach assumes that the intensity value of each pixel hardly varies across images of the muscle at rest, but when a twitch is evoked there is a local, transient variation in the intensity value of the pixels in the area of the image the muscle twitch occurred. The intensity value of each pixel in the first 500 images is therefore used to construct a Gaussian mixture model (KaewTraKulPong and Bowden, 2002). Here X distributions were used. The distributions are weighted, based on the proportion of the image sequence its intensities occur. Intensities in more highly weighted distributions occur more commonly (i.e. when the muscle is at rest), while intensities in less weighted distributions occur less commonly (i.e. only during an evoked twitch). The mixture model is then used to categorise pixels in all subsequent images (>500) as either background or foreground, with the mixture model updating to adapt to any repetitive changes in pixel intensity value (e.g. related to breathing patterns). Images recorded during an evoked twitch will, therefore, contain dense clusters of foreground pixels located in the area the muscle tissue displacement occurred. Connective components were used to analyse the density of foreground pixels in each image, and more sparsely distributed foreground pixels (i.e. resulting from noise) were discarded (Stauffer and Grimson, 1999). The final result was, therefore, a 1-D signal of the number of foreground objects in each image frame, with greater numbers of foreground objects indicating large muscle tissue displacement.

## Results

### Resting Motor Thresholds (RMT)

The results are presented in Figure 2. We report that in BI, FDI and TA, the MEP amplitude, as measured by sEMG, and the tissue displacement, as measured by pUS, increases with stimulation intensity, after reaching a resting motor threshold intensity. RMT intensity varies amongst individuals. Average RMT intensities are lowest in FDI, where most individuals respond with changes in amplitude at 30-50% MSO. Individual BI RMT ranges from as low as 40% MSO and in most cases reach threshold intensity by 70% MSO. RMTs are highest in TA. With the exception of one individual, who’s threshold is below 60% MSO, other individuals had an RMT of 70-90% MSO in TA. RMT intensity measurements appear consistent across MEP and pUS methods on an individual and on a group level.

**Figure 2.**
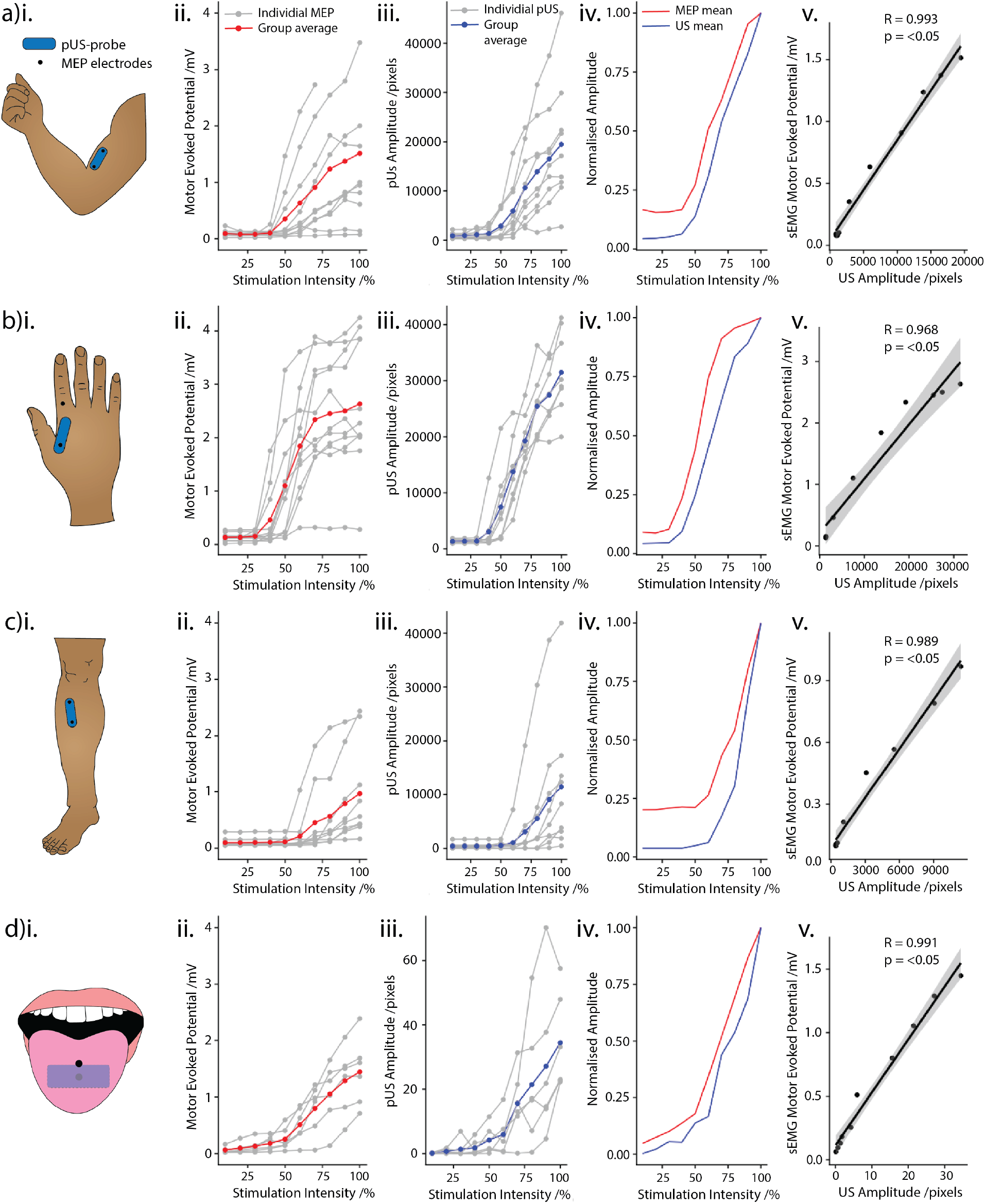
**i.** Cartoon of muscle with the ultrasound probe (blue) and EMG electrode (black) placement on the **(A)** biceps (BI), **(B)** first dorsal interosseous (FDI), **(C)** tibialis anterior (TA), (D) tongue (TO). **ii.** MEP recruitment curve-MEP amplitudes, as measured by EMG of muscle, at TMS stimulation intensities 10%-100% MSO at an individual level. The red line indicates mean values. **iii.** pUS recruitment curve-pUS displacement amplitude*, as measured by pUS of the muscle, at TMS stimulation intensities 10%-100% MSO at an individual level. The blue line indicates mean values. **iv.** Normalised mean amplitudes of TMS-evoked muscle contraction using EMG (red) and US (blue). **v.** Scatterplot indicates a very strong correlation of TMS-evoked muscle contraction using EMG and US. Grey shadow indicates a 95% confidence level interval for predictions from a linear model.

### Amplitude Pattern

Upon visual inspection of Fig 2(v), changes in MEP amplitude and pUS tissue displacement are closely aligned, following a similar sigmoidal pattern in BI, FDI, TA and TO with increasing TMS stimulation intensity. Of interest, the MEP amplitude curve flattens out more towards the high stimulation intensities, whereas this effect is not apparent in the pUS tissue displacement curves. This is particularly highlighted in FDI and BI muscles. All tested muscles show a very high correlation in the changes in amplitudes with respect to stimulation intensity, depicted by sEMG and pUS (TA, R=0.993, p=<0.05; FDI, R=0.968, p=0.05; TA, R=0.989, p=<0.05; TO, R=0.991, p=<0.05).

## Discussion

In this study, we demonstrate that recruitment curves for TMS evoked pUS muscle contraction strongly relates to sEMG. Importantly, the motor evoked threshold defined by pUS is virtually the same as that defined by sEMG. This study shows that pUS is feasible in muscles that are difficult to measure using sEMG. While applications of pUS could be broad, it could be applied to examine the tongue in stroke or MND. This may be particularly useful in clinical studies involving the tongue when repeated measures are required and sEMG may be considered unacceptable. In other words, this work provides a novel alternative to sEMG to measure TMS evoked muscle contraction.

This is the first report of TMS-pUS that we are aware of in the literature. The close relationship of sEMG output and pUS supports the use of pUS in TMS studies. Future experiments need to explore the relationship of more complex TMS protocols, such as short-interval intracortical inhibition (SICI), although we expect the relationship to be conserved. Adapting pUS for use in these types of protocols will enable us to probe excitatory and inhibitory corticomotoneuronal integrity in tracts corresponding to a broader range of muscles. For example, currently, there are very few studies that have explored TMS of some important, but difficult-to-access muscles such as the tongue.

TMS-pUS may be particularly useful in clinical disorders that affect the tongue, such as MND and stroke. There are very few studies that have explored TMS of the tongue, and none that have demonstrated. We suspect this is largely caused by the difficulty in probing this region using sEMG electrodes. MND patients often experience bulbar symptoms and monitoring its progression is currently left to clinical assessments. Understanding the interaction of excitatory and inhibitory networks in the corticomotoneuronal pathways corresponding to bulbar regions may be crucial in the understanding of MND neurophysiology. In stroke, TMS-pUS could allow studies to explore the longitudinal cortical reorganisation that occurs within the tongue region mirror that approach applied to the hand (Sharma and Cohen, 2012).

Our findings suggest that pUS is a real alternative to sEMG. The main advantages for using pUS, for the patient, is that unlike sEMG, setting up pUS to view the tongue is painless, comfortable and non-invasive. We found that tongue sEMG electrodes were not only uncomfortable for our participants, making them more likely to move their tongue during the study, but that throughout the TMS session, saliva builds up causing issues with electrode adhesion and sEMG signal. In many conditions swallowing can lead to excess saliva, which would make the use of electrodes even more challenging. pUS is a good solution to removing discomfort from TMS experiments in the tongue.

There are technical challenges and limitations with TMS-pUS. Ultrasound processing methods are currently carried out offline. This presents challenges when determining the resting motor threshold using pUS alone. In TMS experiments in which RMT is used as a reference point for inhibitory or excitatory protocols, such as short-interval intracortical inhibition (SICI) or short-interval intracortical facilitation (SICF) amongst others, RMT is measured at the start of a testing session. In order for TMS-pUS to be implemented for use instead of sEMG in such protocols, either a fast-offline pUS processing method, or an online pUS processing method must be developed. Secondly, the pUS probe is a manual device in which it may be difficult to achieve reproducible views of a muscle, simply due to area, angle of placement, the pressure applied to the probe or quantity of gel used. Additionally, TMS-pUS only samples one plane of muscles at one time, whereas sEMG can sample multiple superficial muscles.

In summary, this study suggests that TMS-pUS is a novel technique which closely mirrors the sEMG signal. Future work needs to establish its use in paired-pulse and repetitive TMS paradigms. While the off-line processing may limit the use of TMS-pUS in some studies, this is surmountable. Importantly, there are a number of potential applications of TMS-pUS in stroke and neuromuscular diseases.

## ACKNOWLEDGMENTS

We thank Paul Hammond for help with equipment design.

## References

Arts, I.M.P., van Rooij, F.G., Overeem, S., Pillen, S., Janssen, H.M.H.A., Schelhaas, H.J., Zwarts, M.J., 2008. Quantitative Muscle Ultrasonography in Amyotrophic Lateral Sclerosis. Ultrasound in Medicine & Biology 34, 354–361. https://doi.org/10.1016/j.ultrasmedbio.2007.08.013

Bibbings, K., Harding, P.J., Loram, I.D., Combes, N., Hodson-Tole, E.F., 2019. Foreground Detection Analysis of Ultrasound Image Sequences Identifies Markers of Motor Neurone Disease across Diagnostically Relevant Skeletal Muscles. Ultrasound in Medicine & Biology 45, 1164–1175. https://doi.org/10.1016/j.ultrasmedbio.2019.01.018

Darby, J., Hodson-Tole, E.F., Costen, N., Loram, I.D., 2012. Automated regional analysis of B-mode ultrasound images of skeletal muscle movement. Journal of Applied Physiology 112, 313–327. https://doi.org/10.1152/japplphysiol.00701.2011

Lucas, B.D., Kanade, T., n.d. An Iterative Image Registration Technique with an Application to Stereo Vision 9.

Gubbay, SS., Kahana, E., Zilber, N., Cooper, G., Pintov, S., Leibowitz, Y., 1985. Amyotrophic lateral sclerosis. A study of its presentation and prognosis. J Neurol 232, 295–300. https://doi.org/10.1007/BF00313868

Harding, P.J., Loram, I.D., Combes, N., Hodson-Tole, E.F., 2016. Ultrasound-Based Detection of Fasciculations in Healthy and Diseased Muscles. IEEE Trans. Biomed. Eng. 63, 512–518. https://doi.org/10.1109/TBME.2015.2465168

Heckmatt, J.Z., Leeman, S., Dubowitz, V., 1982. Ultrasound imaging in the diagnosis of muscle disease. The Journal of Pediatrics 101, 656–660. https://doi.org/10.1016/S0022-3476(82)80286-2

KaewTraKulPong, P., Bowden, R., 2002. An Improved Adaptive Background Mixture Model for Real-time Tracking with Shadow Detection, in: Remagnino, P., Jones, G.A., Paragios, N., Regazzoni, C.S. (Eds.), Video-Based Surveillance Systems. Springer US, Boston, MA, pp. 135–144. https://doi.org/10.1007/978-1-4615-0913-4_11

Leon-Sarmiento, F.E., Rizzo-Sierra, C.V., Bayona, E.A., Bayona-Prieto, J., Doty, R.L., Bara-Jimenez, W., 2013. Novel Mechanisms Underlying Inhibitory and Facilitatory Transcranial Magnetic Stimulation Abnormalities in Parkinson’s Disease. Archives of Medical Research 44, 221–228. https://doi.org/10.1016/j.arcmed.2013.03.003

Miguez, D., Hodson-Tole, E.F., Loram, I., Harding, P.J., 2017. A technical note on variable inter-frame interval as a cause of non-physiological experimental artefacts in ultrasound. R. Soc. open sci. 4, 170245. https://doi.org/10.1098/rsos.170245

Pascual-Leone, A., Tormos, J.M., Keenan, J., Tarazona, F., Cañete, C., Catalá, M.D., 1998. Study and Modulation of Human Cortical Excitability With Transcranial Magnetic Stimulation: Journal of Clinical Neurophysiology 15, 333–343. https://doi.org/10.1097/00004691-199807000-00005

Reimers, C.D., Ziemann, U., Scheel, A., Rieckmann, P., 1996. Fasciculations: Clinical, electromyographic, and ultrasonographic assessment. J Neurol 243, 579–584. https://doi.org/10.1007/BF00900945

Sharma, N., Cohen, L.G., 2012. Recovery of motor function after stroke. Dev. Psychobiol. 54, 254–262. https://doi.org/10.1002/dev.20508

Stauffer, C., Grimson, W.E.L., 1999. Adaptive background mixture models for real-time tracking, in: Proceedings. 1999 IEEE Computer Society Conference on Computer Vision and Pattern Recognition (Cat. No PR00149). Presented at the Proceedings. 1999 IEEE Computer Society Conference on Computer Vision and Pattern Recognition, IEEE Comput. Soc, Fort Collins, CO, USA, pp. 246–252. https://doi.org/10.1109/CVPR.1999.784637

Terao, Y., Ugawa, Y., 2002. Basic Mechanisms of TMS: Journal of Clinical Neurophysiology 19, 322–343. https://doi.org/10.1097/00004691-200208000-00006

Vucic, S., Kiernan, M.C., 2017. Transcranial Magnetic Stimulation for the Assessment of Neurodegenerative Disease. Neurotherapeutics 14, 91–106. https://doi.org/10.1007/s13311-016-0487-6

Vucic, S., Ziemann, U., Eisen, A., Hallett, M., Kiernan, M.C., 2013. Transcranial magnetic stimulation and amyotrophic lateral sclerosis: pathophysiological insights. J Neurol Neurosurg Psychiatry 84, 1161–1170. https://doi.org/10.1136/jnnp-2012-304019

Walker, F.O., Donofrio, P.D., Harpold, G.J., Ferrell, W.G., 1990. Sonographic imaging of muscle contraction and fasciculations: A correlation with electromyography. Muscle Nerve 13, 33–39. https://doi.org/10.1002/mus.880130108

